# Epigenetic rewriting at centromeric DNA repeats leads to increased chromatin accessibility and chromosomal instability

**DOI:** 10.1101/2021.02.22.432244

**Authors:** Sheldon Decombe, François Loll, Laura Caccianini, Kévin Affannoukoué, Ignacio Izeddin, Julien Mozziconacci, Christophe Escudé, Judith Lopes

## Abstract

Centromeric regions of human chromosomes contain large numbers of tandemly repeated α-satellite sequences. These sequences are covered with constitutive heterochromatin which is enriched in trimethylation of histone H3 on lysine 9 (H3K9me3). Although well studied using artificial chromosomes and global perturbations, the contribution of this epigenetic mark to chromatin structure and genome stability remains poorly known in a more natural context. Using transcriptional activator-like effectors (TALEs) fused to a histone lysine demethylase (JMJD2B), we were able to reduce the level of H3K9me3 on the α-satellites repeats of human chromosome 7. We show that the removal of H3K9me3 affects chromatin structure by increasing the accessibility of DNA repeats to the TALE protein. Tethering TALE-demethylase to centromeric repeats impairs the recruitment of HP1α and proteins of Chromosomal Passenger Complex (CPC) on this specific centromere. Finally, the epigenetic re-writing by the TALE-JMJD2B affects specifically the stability of chromosome 7 upon mitosis, highlighting the importance of H3K9me3 in centromere integrity and chromosome stability, mediated by the recruitment of HP1α and the CPC.

## Introduction

The centromere plays a crucial role in the life cycle of cells, ensuring the faithful transmission of genetic information during cell division. Originally defined as the primary constriction of mitotic chromosomes, it is the structure on which the kinetochore assembles, allowing the cell to split its genetic material into two identical halves (Pluta *et al*, 1990). Centromeres are found in all eukaryote species and some centromere-like structures even exist in archaea and bacteria (Gordon & Wright, 2000; Schumacher *et al*, 2015).

Most eukaryotes possess a regional centromere composed of tandemly repeated sequences called satellite DNA. In humans, the main satellite DNA, called α-satellite, is made of 171 bp monomers that can adopt complex organizational patterns called higher-order repeats (HOR) (Hayden, 2012; Fukagawa & Earnshaw, 2014; McNulty & Sullivan, 2018), but monomer length and composition can differ a lot in other species. Despite this variability, all centromeres share a similar chromatin organization. The centromere is defined by the presence of a histone H3 variant called CENP-A (in blue on Fig 1A). This protein acts by recruiting the constitutive centromere-associated network (CCAN) that will serve as a kinetochore assembly platform during mitosis (Stellfox *et al*, 2013; Sharma *et al*, 2019). This domain is usually surrounded by a compact chromatin domain that is enriched in a specific epigenetic mark, the tri-methylation of lysine 9 on histone H3 (H3K9me3, in red on Fig 1A) that extends on several Mbp, over arrays of α-satellites and including other satellite DNA families (Nishibuchi & Déjardin, 2017). This epigenetic mark can be bound by heterochromatin protein 1 (HP1, in purple on Fig 1A) and these two ingredients are the hallmark of the so-called constitutive heterochromatin (Sales-Gil & Vagnarelli, 2020). HP1 is a family of three proteins (α, β, γ) which can bind H3K9me3 thanks to a chromodomain. HP1 mediates the recruitment of SUV39H1 methyltransferase that catalyses the trimethylation of H3K9 which leads to the spreading of the H3K9me3 mark (Mozzetta *et al*, 2015). HP1 can bridge chromatin segments and form liquid droplets *in vitro* (Larson *et al*, 2017; Strom *et al*, 2017). Whether or not this phenomenon of liquid-liquid phase separation is responsible for the formation of compact heterochromatin domains *in vivo* is still debated (Erdel *et al*, 2020). This last domain is called pericentromere while the CENP-A enriched domain represents the core centromere. This centromere chromatin organization is conserved across the eukaryota, while the DNA sequence is not, which points to an essential functional role of this dual centromeric chromatin organization. Several such roles have been proposed ranging from the faithful segregation of chromosomes during mitosis to the regulation of gene expression (Becker *et al*, 2016; Janssen *et al*, 2018; Ninova *et al*, 2019). While the importance of the CENP-A protein for centromere function is largely documented (Müller & Almouzni, 2017; Sharma *et al*, 2019; Mahlke & Nechemia-Arbely, 2020), much less is known regarding the contribution of the pericentromere region and its associated H3K9me3 mark.

**Figure 1:**
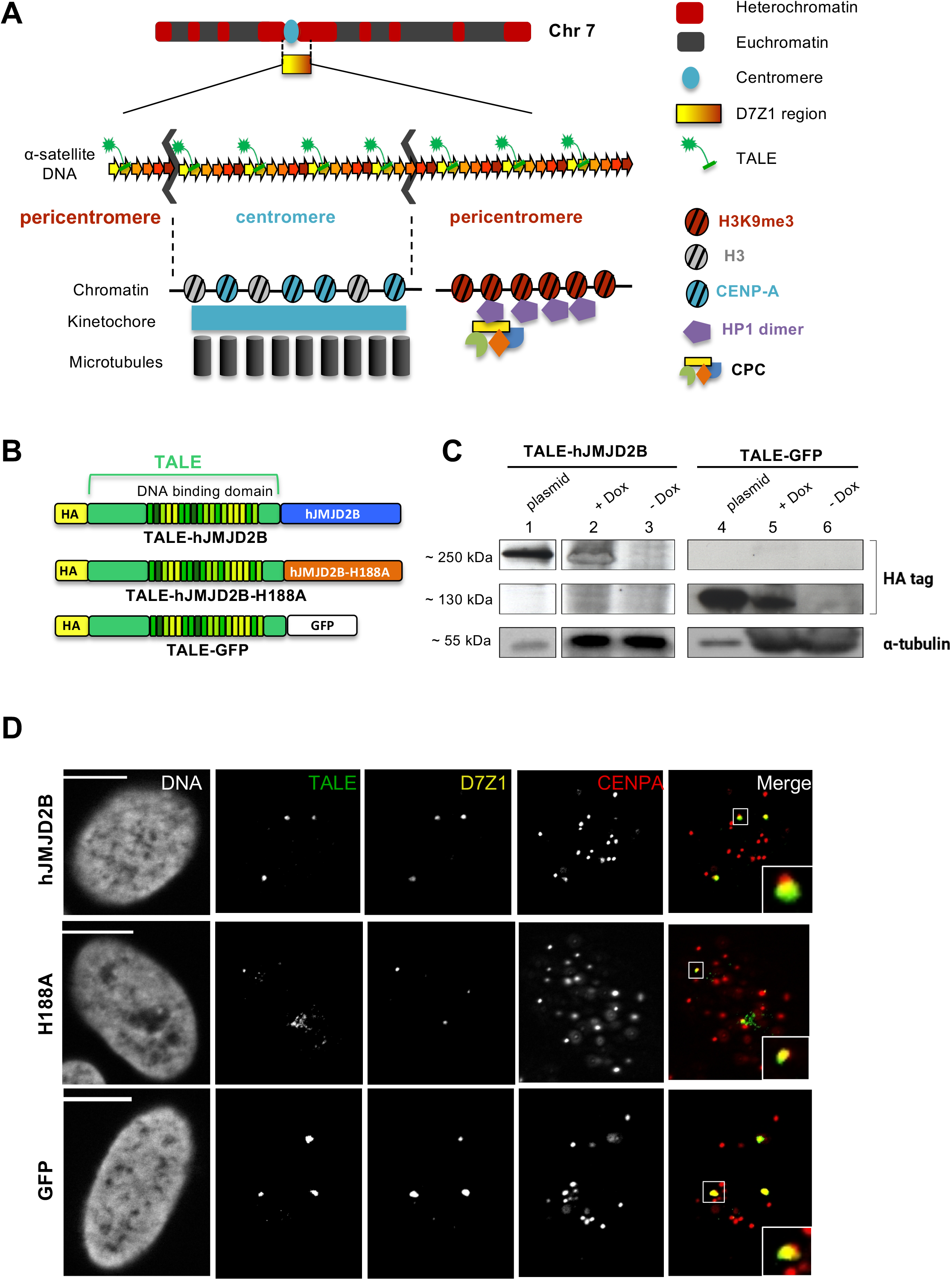
Overview of the system. **(A)** Sketch representing the chromosome 7 with its centromere. Tandem α-satellite repeats of D7Z1 region are represented with the binding of the TALE-effector fusion protein. Chromatin organisation with CENP-A and kinetochore assembly with microtubule attachments, characterizes the core centromere during mitosis. The H3K9me3 epigenetic mark and HP1 protein are the hallmark of the pericentromeric heterochromatin. The CPC, interacting with HP1, is essential to correct erroneous microtubule attachments. **(B)** Sketch of the different constructs of TALE fusion proteins used in this study. **(C)** Western blot against HA revealing TALE-demethylase (~250 kDa), TALE-GFP (~130kDa) and α-tubulin (loading control). The lanes 1 and 4 correspond to transient transfection of cells with the plasmid coding for TALE-demethylase and TALE-GFP, respectively. The + Dox (lanes 2 and 5) and - Dox (lanes 3 and 6) correspond to stable cell lines with or without doxycycline induction of the TALE fusion proteins. Due to the different levels of TALE expression between the two systems (transfection and induction), 1.5 μg of total proteins were loaded for transient transfections while 180 μg were used for the inducible cell lines. **(D)** U2OS cells expressing either the TALE-demethylase (top), its point mutant (middle) or the TALE-GFP (bottom). Cells were stained by IF-FISH, TALE proteins are visualized using an anti-HA antibody, CENP-A using an anti-CENP-A antibody and the D7Z1 region using a specific probe generated at the lab. DNA was stained using Hoechst. Scale bar, 10 μm.

Using the histone lysine methyltransferase *Suv39h double null* mice, Peters *et al*. (2001) reported a reduced pericentromeric H3K9 methylation associated with an increased chromosomal instability. In a similar situation in mouse IMEF cells, McManus *et al*. (2006) also observed mitotic defects. Therefore, preventing H3K9 trimethylation seems to result in defects in the preservation of genome stability. Another approach that can be used to reveal the importance of the H3K9me3 mark consists in removing this mark using a H3K9-specific histone lysine demethylases (Dimitrova *et al*, 2015). In breast tumors exhibiting an over-expression of the H3K9-specific histone lysine demethylase hJMJD2B, Slee *et al.* (2012) observed an increased aneuploidy and chromosomal instabilities. An interesting approach by Molina *et al*. (2016) used another histone lysine demethylase, hJMJD2D, to remove H3K9me3 which resulted in increased mitotic defects and in perturbation of the chromosomal passenger complex (CPC) localization to inner centromeres. During mitosis, the CPC is required to correct erroneous microtubule attachments through the action of Aurora B (Carmena *et al*, 2012; Funabiki, 2019). In this last approach, the activity of the histone demethylase was targeted to heterochromatin by fusing the enzyme to the N terminal domain of SUV39H1 which targets the H3K9me3 mark.

In all aforementioned studies, the removal of the H3K9me3 mark is expected to affect all heterochromatic regions of the genome. Targeted modifications of epigenetic marks to pericentromeric heterochromatin have up to now only been obtained using an artificial system in which a synthetic array of α-satellite DNA of several tens of kb can mimic *de novo* centromere assembly and give rise to the formation of a human artificial chromosome (HAC) (Ohzeki *et al*, 2019; Martins *et al*, 2020). The authors showed that long-term tethering of histone lysine demethylase to HAC, reduced centromere protein levels and triggered HAC mis-segregation, maybe resulting from a decrease of CENP-A level below a critical threshold.

Here, we propose a novel approach, in which the demethylation of H3K9 is targeted to the centromeric region of a specific human chromosome. Although α-satellite monomers usually share at least 60% identity, chromosome specific variations in the composition and organization of monomers have been described (Willard, 1985; McNulty & Sullivan, 2018). Exploiting these variations, we designed a DNA sequence binding domain based on the transcription activator-like effector (TALE) system (Moscou & Bogdanove, 2009; Voytas & Joung, 2009; Jankele & Svoboda, 2014), that specifically targets the D7Z1 array of α-satellite repeats known to be present in the centromeric region of human chromosome 7. We observed that the TALE-fusion protein was specifically recruited at the centromeres of chromosomes 7 and that the targeting of the active histone demethylase resulted in a local decrease of H3K9me3. We determined the effect of this H3K9me3 loss on chromatin organization using both epifluorescence microscopy and super-resolution imaging. Monitoring the corresponding TALE signal also suggests a better accessibility of DNA upon H3K9me3 removal. We further showed that recruitment of the HP1 protein was reduced, inducing in turn a decrease of the chromosomal passenger complex (CPC) at the TALE foci. Finally, we demonstrated that the targeting of the TALE-demethylase to the pericentromeric heterochromatin of chromosome 7 affects the stability of this specific chromosome upon mitosis. This experimental system opens interesting possibilities for further studies regarding the role of the H3K9me3 mark.

## Results

### Targeting α-satellite repeats from a specific human chromosome

Our aim was to modify the centromeric chromatin on a specific pair of chromosomes by fusing the histone lysine demethylase hJMJD2B to a DNA binding domain of a TAL effector. This domain was designed to bind an 18 bp motif that is present in the D7Z1 array of α-satellite repeats known to be present in the centromeric region of human chromosome 7 (Fig 1A). According to the models for α-satellite genome content that have been integrated in the h38 version of the human genome assembly, the D7Z1 region is made of more than 2500 repeats of an HOR made of 6 α-satellite monomers, covering about 2.6 Mbp and contains at least 2200 target sequences for the TALE. This region is flanked on the short arm side by another array of repeats called D7Z2 that is much shorter (about 250 Kbp). ChIP experiments have shown that the centromere of chromosome 7 is located on the D7Z1 array of repeats (Slee *et al*, 2012; Hayden *et al*, 2013). Nevertheless, because D7Z1 extends over the core centromere, we expect the TALE to bind the D7Z1 repeats at both the centromere and the pericentromeric region of chromosome 7 (see below).

As controls, we also fused the TALE to the catalytically inactive histone lysine demethylase (TALE-hJMJD2B-H188A) (Whetstine *et al*, 2006) or to a GFP (TALE-GFP). A HA tag was fused to the N-terminal part of each TALE for visualization using a specific antibody (Fig 1B). These constructs were expressed in U2OS cells by transient transfection of the corresponding plasmid. We also established inducible U2OS cell lines expressing either the TALE-demethylase or TALE-GFP in response to doxycycline. This inducible system was developed to determine the chromosome 7 instability in a large number of cells after mitosis. The expression of our TALE constructs in both systems was confirmed by western blot (Fig 1C). The expression of the TALE-hJMJD2B and of the TALE-GFP upon transient transfection of U2OS (lanes 1 and 4 in Fig 1C) is much higher than the expression of these proteins in the inducible U2OS cell lines after doxycycline treatment. This difference of expression probably results from a higher number of plasmid copies introduced in the nucleus after transient transfection of the cells in comparison with the number of copies stably integrated in the genome of the inducible U2OS cell lines.

We first wanted to demonstrate the specific binding of our TALE targeting the α-satellites of chromosome 7. For that, we transiently transfected U2OS cells with plasmids expressing the different TALE fusion proteins and fixed the cells after 24 hours. We then performed immunofluorescence coupled to fluorescent *in situ* hybridization (IF-FISH) to investigate the localization of the TALE proteins (revealed by an anti-HA antibody) with respect with the centromeric region of chromosome 7 (labelled with a probe targeting the D7Z1 repeats) and the core centromere (visualized by an anti-CENP-A antibody) (Fig 1D). Although the images sometimes showed a faint staining of the nucleolus, four TALE foci were observed in each nucleus, overlapping with the D7Z1 signals. This number is in accordance with the fact that U2OS cells are mostly tetraploids. There was usually one CENP-A foci overlapping with the TALE-D7Z1 foci and the CENP-A foci was slightly smaller than the TALE-D7Z1 signals (see inset in Fig 1D). These results indicate that targeting of the centromeres from chromosomes 7 was achieved and suggest that the binding overlaps a part of the core centromere and the adjacent pericentromeric region.

### TALE protein fused to histone demethylase specifically removes H3K9me3 from chromosome 7 centromere

Having validated the targeting activity of our construct, we next tested the efficiency of the demethylase activity brought by the TALE-hJMJD2B. To do so, the labelling of H3K9me3 revealed by immunofluorescence was quantified in cells either transfected with the active TALE-hJMJD2B or with the inactive TALE-hJMJD2B-H188A. For each condition, the 3D fluorescence images of hundreds of nuclei were analyzed with the software Tools for Analysis of Nuclear Genome Organization (TANGO) (Ollion *et al*, 2013). This tool was developed for quantitative study of large sets of images and statistical processing with R. Quantification of H3K9 tri-methylation levels was performed over the detected TALE foci in each nucleus and the signal was normalized by taking into account the signal present in the whole nucleus. Our experiments revealed a statistically significant loss of H3K9me3 of 19.6% (p<0.0001, Wilcoxon-Mann-Whitney test) at the binding sites of the TALE-demethylase, in comparison with the TALE-demethylase inactive (Fig 2A). Our control using an inactive demethylase ensures that this loss is mediated by the histone lysine demethylase activity and is not merely a consequence of the binding of the TALE protein.

**Figure 2:**
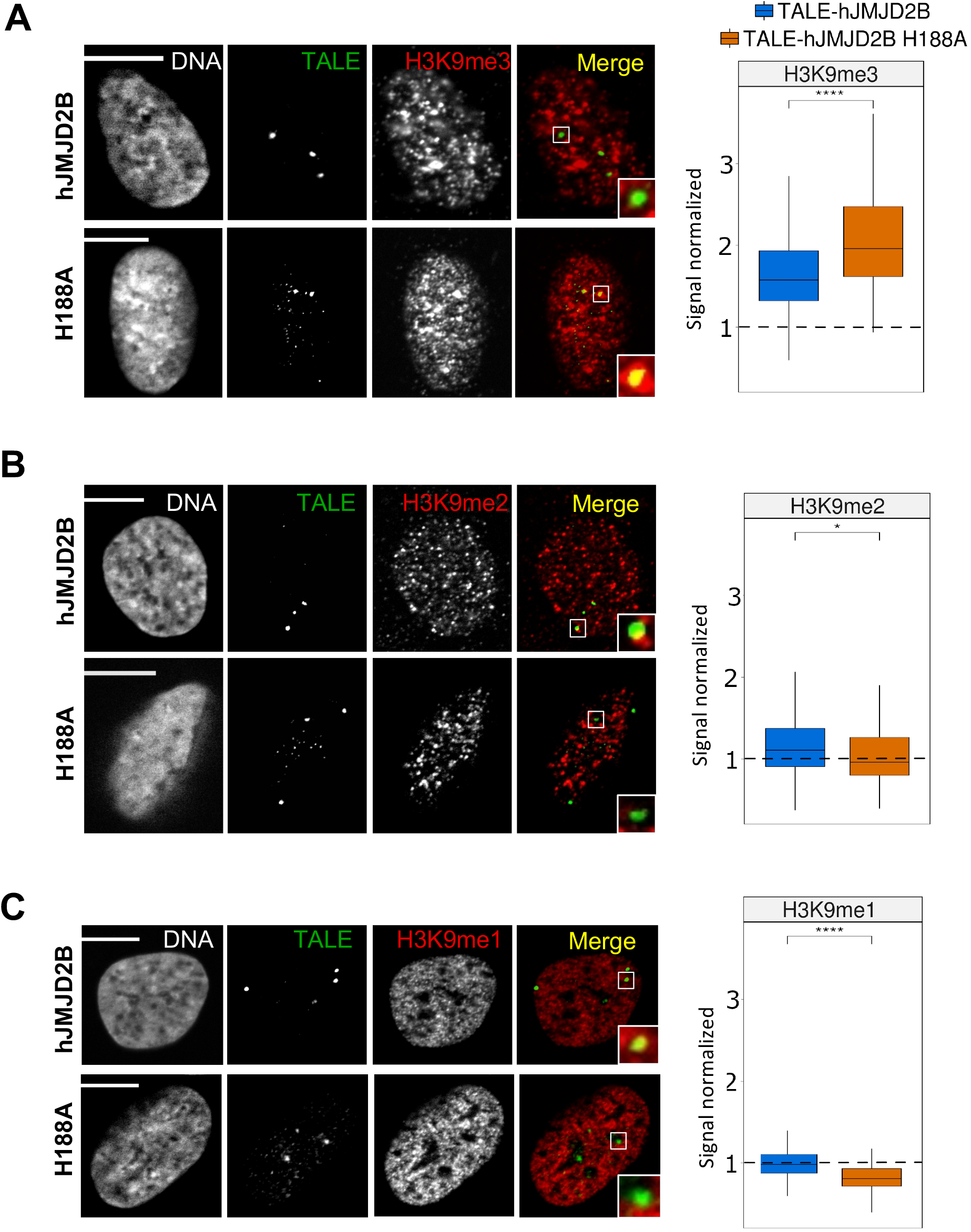
Effects of TALE-demethylase expression on H3K9 methylation. U2OS cells expressing the TALE-demethylase (top) or the catalytically inactive mutant (bottom). DNA is visualized using Hoechst, the TALE (shown in green) is revealed with an anti-HA antibody and H3K9me3 **(A)**, H3K9me2 **(B)** or H3K9me1 **(C)** (shown in red) is revealed with a specific antibody. Right panels: boxplots showing the signal normalized at the TALE-demethylase foci (blue) (H3K9me3: n=215 nuclei in 3 experiments, H3K9me2 / H3K9me1: n=128 nuclei in 2 experiments) or at the point mutant (orange) (H3K9me3: n=129 nuclei in 3 experiments, H3K9me2 / H3K9me1: n=123 nuclei in 2 experiments). Signal in the foci is normalized by the global signal in the nucleus. The box represents 50% of data points and the whiskers extend up to 1.5 times the interquartile range, the horizontal bar represents the median. The dashed line represents a signal normalized equal to one. p-values were computed using a Wilcoxon-Mann-Whitney test (*: p < 0.05 / **: p < 0.01 / ***: p < 0.001 / ****: p < 0.0001). Scale bar, 10 μm.

Using the same method, we also measured the levels of H3K9 di- and mono-methylation with specific antibodies. Both levels of H3K9me2 and H3K9me1 increased (by 15.3% and 21.5%, respectively) in response to the removal of one methyl group from H3K9me3 (Fig 2 B-C). We concluded that the recruitment of the TALE-demethylase at the α-satellite repeats on D7Z1 region induced a significant loss of tri-methylation of H3K9 at the targeted sites, associated with an increase of di- and mono-methylation of H3K9.

### Removal of H3K9me3 increases the DNA accessibility but does not change the size of TALE foci

The H3K9me3 epigenetic mark is thought to play a major role in the chromatin accessibility and compaction by recruiting HP1 and the methyltransferases SUV39H and indirectly by favouring the hypoacetylation of histone H3 (Allshire & Madhani, 2018). We thus wondered how the decrease of H3K9me3 triggered by the TALE-demethylase, affects the chromatin accessibility and compaction. We noticed that foci of the inactive TALE-hJMJD2B-H188A seem often less intense that those of the TALE-demethylase, suggesting a higher recruitment of the TALE-demethylase leading to more intense foci.

In order to measure this effect, we estimated the chromatin accessibility of the targeted sites by measuring the labelling intensity of the TALE proteins in these foci. We co-transfected U2OS cells with two distinct plasmids, the first one that expresses either the TALE-hJMJD2B or the TALE-hJMJD2B-H188A (TALEs with HA tag) and the second one that expresses the TALE protein without effector (TALE with FLAG tag) (Fig 3A). Both TALEs target the same sequence on α-satellites of chromosome 7. We detected the presence of the two co-transfected plasmids in U2OS cells by using dedicated antibodies against HA and FLAG tags. We measured the occupancy of the TALE without effector with the FLAG antibody, in the two different cellular contexts (in presence of TALE-demethylase or TALE-hJMJD2B-H188A).

**Figure 3:**
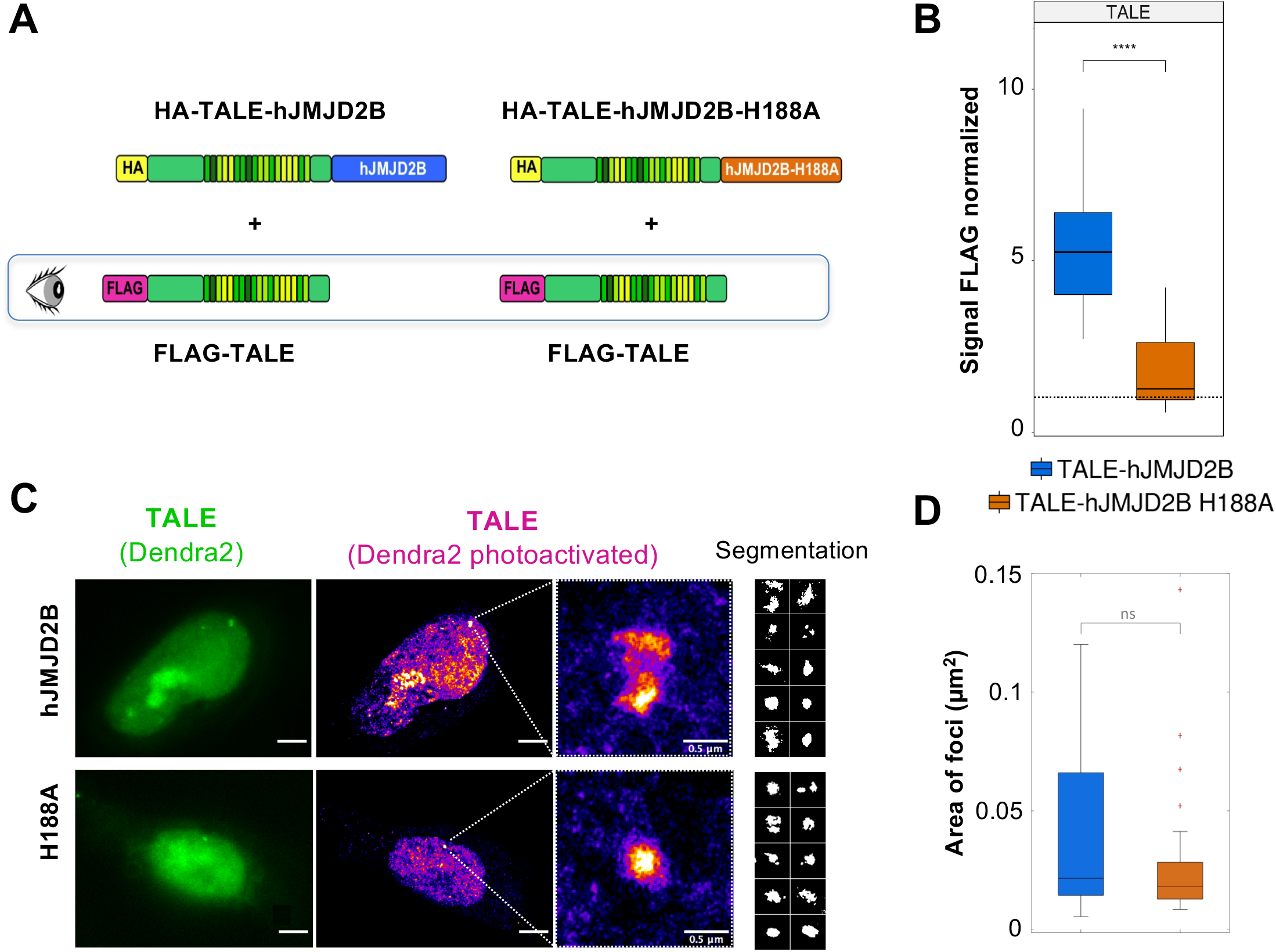
DNA accessibility and local compaction of α-satellite target sites. **(A)** Sketch representing the co-transfection of two TALEs targeting the D7Z1 α-satellite repeats. The FLAG-TALE (without effector) is transfected with the TALE-hJMJD2B or with the TALE-hJMJD2B-H188A. Only the FLAG signal is quantified in the cells expressing the two constructs. **(B)** Boxplots showing the FLAG signal normalized in the cells co-expressing the TALE-hJMJD2B (blue) (n=50 nuclei) or the point mutant (orange) (n=37 nuclei). The dashed line represents a signal normalized equal to one. **(C)** Measure of the TALE foci area by super-resolution microscopy (PALM). U2OS cells were transfected with the TALE-hJMJD2B-Dendra2 (top) or the TALE-hJMJD2B-H188A-Dendra2 (bottom). Cells are visualized in classical epifluorescence after an excitation at 488 nm (left, green), and super-resolution images reconstructed after photoactivation (405 nm) and excitation (561 nm) of Dendra2 single-molecule events (middle). Zoom of foci are presented on the right, as well as some examples of segmented foci. Scale bar, 5 μm and 0.5 μm for the right panel, segmented clusters images are 1 μm x 1 μm. **(D)** Boxplots representing the measure of the area of foci segmented of the TALE-hJMJD2B-Dendra2 (blue) (n= 27 foci) or the TALE-hJMJD2B-H188A-Dendra2 (orange) (n= 36 foci), with median values of 0.022 μm^2^ and 0.018 μm^2^ for the TALE-demethylase and TALE-inactive, respectively.

The signal of the FLAG antibody is much higher in the context of H3K9me3 decrease, with a normalized signal of 5.4 in the presence of the TALE-demethylase and 1.8 in presence of the TALE-hJMJD2B-H188A (Fig 3B). The 3-fold increase of the FLAG signal suggests that the partial removal of H3K9me3 leads to a higher accessibility of the TALE to its binding sites.

We next asked whether the increased accessibility of the targeted sites is associated with change in chromatin compaction that would result in an increase of the TALEs foci volume. We turned to super-resolution microscopy and used photo-activated localization microscopy (PALM) in order to obtain images with a resolution beyond the diffraction limit that would allow us to monitor slight changes in the foci volumes. We fused the Dendra2 protein to the C-terminal part of the TALE-demethylase and TALE-hJMJD2B-H188A. Dendra2 is a photoconvertible green-to-red fluorescent protein, commonly used for single-molecule microscopy super-resolution imaging (SMLM). We identified several foci of TALE-demethylase and TALE-hJMJD2B-H188A and measured the area of these foci (Fig 3C). While demethylated foci showed a higher size variability, we did not detect a significative change of the average size of the foci (p=0.13, Wilcoxon rank sum test, Fig 3D). We could not rule out that the higher size variability of demethylated foci could be explained by the copy number variation of plasmids expressing the TALE-demethylase in the cells.

Taken together, the results obtained with the two approaches above, reveal that the higher accessibility of the α-satellites DNA repeats induced by the removal of H3K9me3 is not associated with a visible decompaction of the chromatin.

### Removal of pericentromeric H3K9me3 results in chromosome specific instability

To test whether the H3K9me3 had a role in ensuring the proper segregation of sister chromatids, we measured the impact of H3K9me3 loss, brought by the TALE-demethylase, on the stability of chromosome 7 and chromosome 11 used as a control. We grew the two doxycycline inducible cell lines expressing either the TALE-demethylase or the TALE-GFP for 48h with or without doxycycline and determined the number of chromosome 7 and 11 in each nucleus. FISH images in which the centromeres of chromosomes 7 and 11 were labelled with centromere specific probes were acquired, covering over a thousand nuclei for each condition. The number of foci was used as a measure of genome stability: the higher the number of nuclei containing over or under 4 chromosomes (U2OS cells being tetraploid for chromosomes 7 and 11), the higher the instability for this chromosome. In our TALE-demethylase cell line, we measured a significant 64% increase of chromosome 7 instability (p<0.01, Chi-squared test), going from 45 nuclei (3.9%) containing an altered number of chromosomes 7 in cells cultivated without doxycycline to 77 nuclei (6.4%) in cells grown with doxycycline, which triggers the expression of the TALE-demethylase (Fig 4 and Table S1). The number of nuclei containing an altered number of chromosomes 11 after 48h of growth went from 27 without doxycycline to 34 with the drug. However, this slight variation proved to be non-significant (p = 0.55). In our TALE-GFP cell line, we did not observe any significant variation between doxycycline-treated and non-treated cells, neither for chromosome 7 nor for chromosome 11 (Fig 4 and Table S1). This result suggests that the binding of the TALE-GFP does not affect the chromosomes segregation during mitosis. We notice a difference in the frequency of instability for chromosome 7 in the TALE-demethylase (3.9%) versus the control TALE-GFP cell line (1.7%) in absence of doxycycline. This could be due to a transcriptional leakage of the TALE-demethylase even though we could not detect it by western blot.

**Figure 4:**
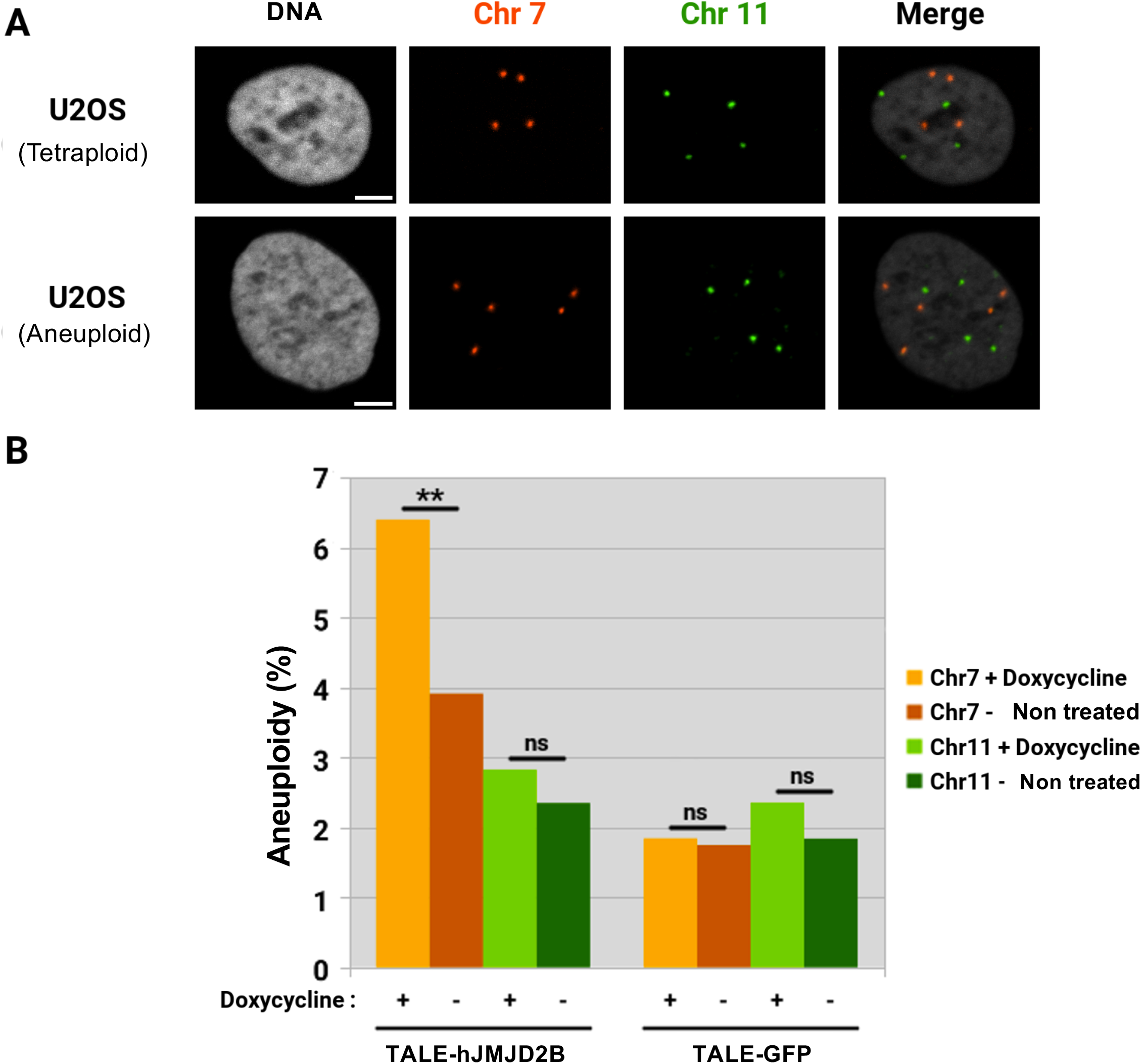
Chromosome instability upon removal of chromosome 7 pericentromeric H3K9me3. **(A**) U2OS cells expressing the inducible TALE-demethylase. DNA is visualized using Hoechst, the α-satellite repeats of chromosome 7 and 11 (shown in orange and green, respectively) are labelled using specific LNA probes. **(B)** Bar plot showing the level of chromosome instability (percentage of nuclei with more or less than 4 chromosomes) in over 1,000 nuclei for chromosome 7 (orange) or chromosome 11 (green) upon expression (light) or not (dark) of the TALE-demethylase (left) or of the TALE-GFP (right). p-values were computed using a Chi-squared test (**: p < 0.01). Scale bar, 5 μm.

We conclude that the loss of H3K9me3 in the pericentromeric region of a chromosome is responsible for mitotic defects that can change the ploidy of this chromosome.

### Partial removal of pericentromeric H3K9me3 is sufficient to affect HP1α and CPC recruitment

Having successfully showed that modifying the epigenetic profile of the centromeric region results in mitotic defects, we wanted to explore the recruitment of some proteins involved in regulation of key mitotic events. The presence of the Chromosomal Passenger Complex (CPC) is essential during interphase and mitosis to correct the chromosome microtubule attachment errors, and activate the spindle assembly checkpoint to insure the proper segregation of chromosomes. This complex is composed of several proteins, Survivin, INCENP, Borealin and its catalytic subunit Aurora B Kinase (Carmena *et al*, 2012). The HP1α protein interacts with INCENP and Borealin and this interaction specifies the CPC localization in the centromere (Ainsztein *et al*, 1998; Liu *et al*, 2014).

In order to link the increased chromosomal instability to a potential change in partners recruited at this specific location, we determined how the decrease of H3K9me3 triggers by the TALE-demethylase affected firstly the HP1 recruitment and secondly the binding of the CPC. We measured the levels of the immunostaining of HP1α with a dedicated antibody at the TALE foci, in cells which expressed the active versus inactive TALE-demethylase. We observed a 22% decrease of HP1α labelling at the binding site of the TALE-demethylase compared to the TALE-inactive (Fig 5A). This result indicates that the loss of H3K9me3 induces a loss of HP1α in the same proportion, underlining the direct interplay between these proteins (Nishibuchi & Nakayama, 2014; Kumar & Kono, 2020).

**Figure 5:**
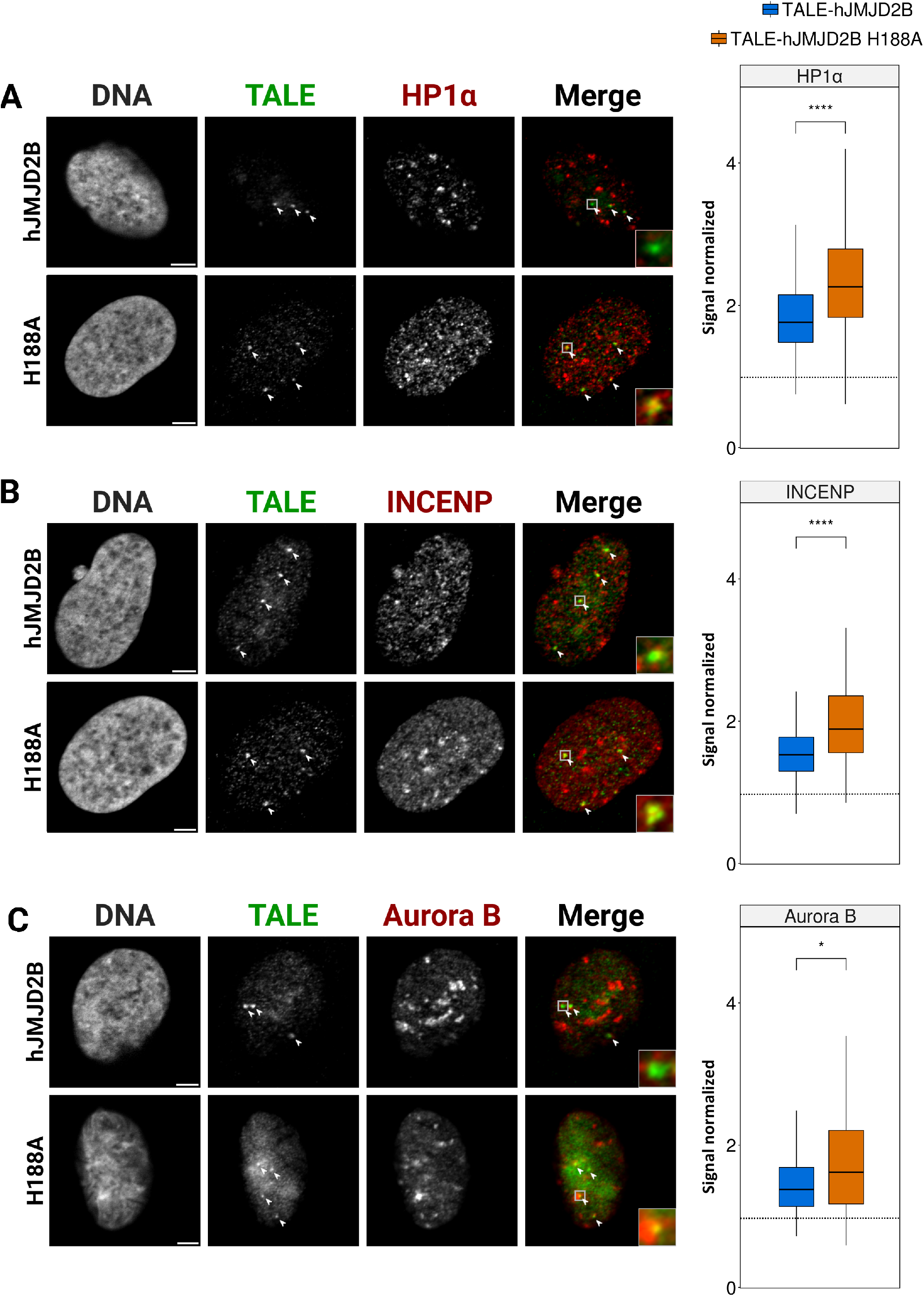
Effects of TALE-demethylase expression on the density of pericentromeric proteins. U2OS cells expressing the TALE-demethylase (top) or the catalytically inactive mutant (bottom). DNA is visualized using Hoechst, the TALE (shown in green) is revealed with an anti-HA antibody and HP1α **(A)**, INCENP **(B)** or Aurora B **(C)** (shown in red) is revealed with a specific antibody. White arrows indicate TALE foci in each cell. Boxplots showing the signal at the TALE-demethylase foci (blue) (HP1α: n=607 nuclei in 5 experiments, INCENP: n=282 nuclei in 2 experiments, Aurora B: n=153 nuclei in 2 experiments) or at the point mutant (orange) (HP1α: n=308 in 5 experiments, INCENP: n=130 in 2 experiments, Aurora B: n=118 in 2 experiments). Signal in the foci is normalized by the global signal in the nucleus for HP1α, INCENP and Aurora B. The box represents 50% of data points and the whiskers extend up to 1.5 times the interquartile range, the horizontal bar represents the median. The dashed line represents a signal normalized equal to one. p-values were computed using a Kruskal-Wallis test (*: p < 0.05 / **: p < 0.01 / ***: p < 0.001 / ****: p < 0.0001). Scale bar, 5 μm.

We then examined the recruitment of INCENP and Aurora B, two proteins of the chromosomal passenger complex. Both for INCENP and Aurora B, we measured a decrease of the signal at the TALE-demethylase foci of 14.9% and 19.1%, respectively, when compared to the point mutant (TALE-hJMJD2B-H188A) (Fig 5B-C). Taken together, these results show that the partial removal of H3K9me3 is sufficient to remove an equivalent amount of HP1α, that in turn affects the recruitment of the CPC at pericentromeres.

## Discussion

Using TALE fused to histone lysine demethylase targeting α-satellites repeats of chromosomes 7, we were able to reduce the level of H3K9me3 in the heterochromatin region of these chromosomes. Our approach has the dual advantage to target endogenous centromeric regions with native chromatin and to target only one specific chromosome so that the effect of the modification induced can be specifically assessed by comparison with other chromosomes. With this new epigenetic tool, we deciphered the consequences of removing the H3K9me3 heterochromatin mark on the centromere of chromosome 7 and assessed the effect of this epigenetic re-writing on chromatin accessibility, chromatin compaction, on centromeric proteins recruitment and on the stability of the chromosomes upon mitosis.

We showed that the targeting of the TALE-demethylase on D7Z1 region induced a significant loss of tri-methylation of H3K9, associated with an increase of di- and mono-methylation of H3K9. This reduced level of H3K9me3 was accompanied by an expected decrease of HP1α and of two major actors of the CPC. These changes in protein composition of the centromeric chromatin were associated with an increase of chromatin accessibility. Chromatin accessibility is usually measured using enzymes that digest, cut or modify accessible DNA and high throughput sequencing (Klemm *et al*, 2019). These techniques are difficult to calibrate in the context of repetitive DNA and they are restricted to a definition on accessibility that rules out potential binding on nucleosomal DNA. It is however now well accepted that some transcription factors can bind DNA wrapped onto nucleosomes (Fernandez Garcia *et al*, 2019). We turned here to another quantification of chromatin accessibility based on microscopy. We used the intensity of labelled TALE targeting sequences found in the foci of interest to assess the local accessibility of the DNA sequences within these foci. Our experiments, carried in two different H3K9me3 chromatin contexts showed that chromatin lacking H3K9me3 is more prone to be bound by sequence specific proteins such as TALEs. Interestingly, we also showed by super-resolution microscopy that this higher DNA accessibility is not associated with a larger size of TALE foci. This result reveals that an increase of DNA accessibility is not always accompanied by a visible decompaction of the chromatin.

We observed that the removal of H3K9me3 was associated with a decrease of INCENP and Aurora B, two members of the CPC. This result is in line with the loss of Survivin (another protein of the CPC) upon removal of H3K9me3 in interphase reported by Molina *et al*. (2016) and suggests that the entire CPC formation could be impaired. The CPC is known to localize to pericentromeres during late S-phase by binding to HP1α, allowing Aurora B to phosphorylate histone H3 at serine 10 (H3S10ph) (Hayashi-Takanaka *et al*, 2009; Carmena *et al*, 2012). This phosphorylation event induces the release of HP1α from chromatin and is thought to be involved in chromosomal condensation at the onset of mitosis (Wei *et al*, 1999; Giet & Glover, 2001; Kruitwagen *et al*, 2015). The transient inhibition of Aurora B activity during interphase has been shown to lead to aberrant chromosome condensation and segregation (Lavoie *et al*, 2004; Petrova *et al*, 2013; Ohishi *et al*, 2014). In a recent study, the authors altered the HP1α dynamics by strongly tethering HP1α to centromeres inducing the CPC recruitment constitutively throughout the cell cycle (Ruppert *et al*, 2018). This tethering caused mitotic delays and high frequency of chromosome segregation errors. In the present work, we complement these studies by showing that the recruitment of the CPC by HP1 is already active in interphase and is impaired by a loss of H3K9me3. Taken together, these results show that the fine tuning of HP1 dynamics is necessary for the correct recruitment and activation of the CPC, an essential factor that controls chromosome segregation during mitosis. Another role of HP1 in mitosis progression is related to the chromoshadow domain of HP1α that supports proper centromeric cohesion by interacting with the Haspin kinase (Yi *et al*, 2018). The double null HP1α and HP1γ cells present indeed prolonged mitosis, increased chromosome mis-segregation and premature chromatid separation.

Looking at the number of chromosomes 7 and 11 after cells underwent mitosis, we were able to observe an increase in mis-segregation for chromosome 7 but not for chromosome 11 on which no epigenetic modification has been targeted. Taken together, these results show that pericentromeric H3K9me3 plays a role in the correct repartition of chromosomes during mitosis by acting as a platform to locate HP1α which, in turn, recruits the CPC. For the first time, we demonstrate the role in genome stability of pericentromeric H3K9me3 in a chromosome-specific manner. Other studies have shown a correlation between H3K9me3 loss and an increase of mis-segregation of chromosomes but they involved a whole H3K9me3 removal on all heterochromatin regions, triggered either by the double inactivation of SUV39H1/2 genes or the over-expression of histone lysine demethylase (Peters *et al*, 2001; McManus *et al*, 2006; Slee *et al*, 2012; Molina *et al*, 2016). Our study shows that the chromosomal instability reported in these studies is not the result of a secondary event like an oncogene activation but rather results from the impaired assembly of the CPC at centromeres.

Our inducible cellular models offer perspectives to decipher the role of H3K9me3 mark in structural aspects and fine regulation of centromeric chromatin (Schalch & Steiner, 2017), as well as, in genome 3D organization and gene regulation where the importance of centromeres is now recognized (Muller *et al*, 2019; Ninova *et al*, 2019).

## Materials and Methods

### Cell culture

U2OS cells were maintained in McCoy’s 5A (1X) + GlutaMAX™ medium supplemented with 10% FBS in a cell culture incubator (37°C in 5% CO_2_). Transfections were performed using the AMAXA™ Cell Line Nucleofector™ Kit V (Lonza): cells were resuspended using 0.05% Trypsin-EDTA(1X) and counted on a Malassez counting chamber, then 10^6^ cells were transfected with 2μg of plasmid. They were put back in culture on a 22×22×0.17mm coverslip in a 6 wells plate for 24h. Stable cell lines were induced with 2 μg/mL doxycycline for 48h prior to analysis.

### Plasmids

The TALE proteins, produced in-house by the TACGENE platform, are composed of the N-Terminal Region (NTR) and the Central Repeat Domain (CRD) with 17.5 repeated motifs binding the following α-satellite sequence from D7Z1 centromeric region: 5’TGCAATTGTCCTCAAATC 3’. In the N-terminal part of the TALE, the NLS from SV40 and three HA tags were added. At the C-terminal part of the TALE was fused either the cDNA of the wild-type histone lysine demethylase hJMJD2B (kindly given by Sophie Beyer), or the hJMJD2B cDNA with the H188A point mutation or a GFP. The H188A mutation was introduced by QuikChange II Site-Directed Mutagenesis kit (Agilent Technologies) with the following primers: 5’GAAGACCACCTTCGCCTGGGCCACCGAGGACATGGACCTGTA3’ and 5’TACAGGTCCATGTCCTCGGTGGCCCAGGCGAAGGTGGTCTTC3’. In transient transfections, constructs were expressed under a CMV promoter.

Cell lines were generated from U2OS cells using the Tet-On^®^ system from Clontech^®^ and following the manufacturer’s instructions. In these stable cell lines, two plasmids were integrated successively. The first expresses the transactivator protein under a CMV promoter and possess a G418 resistance gene for selection. The second contains our TALE fusion protein (TALE-hJMJD2B or TALE-GFP) expressed under the Ptre3g promoter containing seven Tet-O repeats and was co-introduced in cells with a linear puromycin marker for selection. Individual colonies were isolated using 150 μL glass cloning cylinders from Sigma-Aldrich^®^. The list of plasmids we used is presented in Table S2.

### Western blot

Protein samples were separated by SDS-PAGE after 1,5 hours of migration at 150V in Tris-Glycine-SDS buffer 1X. Proteins are transferred onto nitrocellulose membranes (Bio-Rad) by liquid transfer over night at 30V with Tris-Glycine-SDS buffer 1X / EtOH 10%. Membranes are probed with the following antibodies: HA (11867423001, Roche) and α-tubulin (T5168, Sigma-Aldrich). Revelation of the membranes are performed with the ECL Prime Western blotting detection reagent (Amersham).

### Immunofluorescence and FISH

IF and FISH protocols used were the same as described in (Ollion *et al*, 2015b). Briefly, cells were fixed in 4% paraformaldehyde (PFA) in 1X PBS for 10 min and washed three times with PBS. Cells were permeabilized with 0.1% Triton X-100/PBS for 5 min, washed two times with PBS and blocked with 1.5% blocking reagent (Roche Applied Science) at 37°C for 30 min. Primary antibodies were incubated for 1.5 h at room temperature (RT). After three washing with 0.1% Triton X-100/PBS at RT, secondary antibodies were incubated for 1 h at RT in the dark, followed by three washing. For immunofluorescence experiment, DNA staining was performed with 1/5000 dilution of Hoechst 33342 solution (Thermo Scientific) for 10 min. For IF-FISH experiment, a post-fixation step is performed with PFA 2% in 1X PBS for 10 min and washed three times with 0.05% Triton X-100/PBS. Nuclei were permeabilized with 0.5% Triton X-100/PBS for 10 min, rinsed three times with 0.05% Triton X-100/PBS, treated with 0.1N HCl for 1 min, rinsed twice, equilibrated in 2X SSC, and then treated with 2X SSC/50% deionized formamide at room temperature for at least 30 min before hybridization. Oligonucleotide probes were diluted at 0.2 μM in a hybridization solution (2X SSC, 50% deionized formamide, 1X Denhardt solution, 10% dextran sulfate, 0.1% SDS). Hybridization is performed in an *in situ* PCR apparatus (MJ Research) with the following steps: 3 min at 85°C then slowly cooled to 37°C at a rate of 1°C/s and an incubation at 37°C for 2 min. Cells were washed twice in 2X SSC at 63°C. If a secondary antibody is necessary to reveal the probe, cells were incubated in blocking solution (4% BSA, 1X PBS, 0.05% Tween 20) for 30 min at 37°C. Secondary antibodies were diluted in blocking solution and incubated for 1 h at RT in the dark, followed by three washing in 0.05% Tween 20/ 2X SSC. DNA was stained with Hoechst solution, then coverslips were mounted with PPD8 mounting medium (Sigma-Aldrich).

For PALM experiments, IF was performed but the sample were not stained with Hoechst, instead they were stored in 1X PBS at 4°C and protected from light until image acquisitions. At that time, a 1:100 dilution of 0.1 μm TetraSpeck™ microspheres was applied to the samples which were then covered in 50 mM MEA/GLOX buffer and mounted onto a 70 x 24 x 0.17 mm coverslip.

### Antibodies and probes

#### Primary antibodies

Aurora B (ab2254, Abcam), CENP-A (ab13939, Abcam), H3K9me1 (39681, Active Motif), H3K9me2 (05-1249, Millipore), H3K9me3 (07-523, Upstate), HA (ab9111, Abcam), HP1α (MAB3584, Millipore), INCENP (ab36453, Abcam).

#### Secondary antibodies

anti-chicken Alexa Fluor® 488 (703-546-155, Jackson ImmunoResearch), anti-mouse Cyanine 3 (115-167-003, Jackson ImmunoResearch), anti-rabbit Cyanine 3 (111-166-045, Jackson ImmunoResearch), anti-mouse DyLight® 649 (715-495-150, Jackson ImmunoResearch), anti-rabbit Cyanine 5 (111-175-144, Jackson ImmunoResearch), anti-digoxygenin TAMRA (11207750910, Roche) *(used for FISH)*. Primary and secondary antibodies were usually diluted at 1:200.

Centromere-specific oligonucleotide probes were used for the FISH experiments: the a7d probe (5’ A<u>T</u>T<u>G</u>T<u>C</u>C<u>T</u>C<u>A</u>A<u>A</u>T<u>C</u>G<u>C</u>T<u>T</u> 3’) targets the D7Z1 α-satellite repeats while the a11G probe (5’ A<u>G</u>G<u>G</u>T<u>T</u>T<u>C</u>A<u>G</u>A<u>G</u>C<u>T</u>G<u>C</u>T<u>C</u> 3’) targets the D11Z1 repeats (nucleotides underlined indicate LNA bases). The a7d oligonucleotide is coupled with digoxigenin and a11G with AF488 fluorophore. The D7Z1 probe used for IF-FISH was generated by random priming, the PCR product used for the reaction was obtained by amplifying U2OS DNA with the following primers: Fwd 5’ ACGGGGTTTCTTCCTTTCAT / Rev 5’ GCTCTCTCTAAAGCAAGGTTCA.

### Microscopy and 3D image analysis

Classical microscopy imaging was performed with a Zeiss epifluorescence inverted microscope (Axio Observer Z1) equipped with a plan-Apochromat 63×/1.4 numerical aperture oil-immersion objective, as described in (Ollion *et al*, 2015b). Z-stacks consisted of 50–60 sequential slices of 1344 × 1024 pixels (pixel size, 0.1 μm) captured at 0.23 μm intervals by a CCD ORCA-R2 camera (Hamamatsu). For each optical section, images were collected sequentially for each fluorescence channel.

Images were processed using TANGO (v0.93), the nuclei (labelled with Hoechst) were automatically segmented using a home-made 3D watershed-derived algorithm (called Nucleus Edge Detector), which detects nuclear borders using an intensity gradient maxima criterion (Ollion *et al*, 2015a). The TALE foci were segmented using dedicated algorithm (called Spot Segmenter). Quality of segmentation was manually verified in each cell. The different signals corresponding to H3K9me3/me2/me1, HA, FLAG, HP1, Aurora B and INCENP, were measured inside of the segmented structures (the nuclei and the TALE foci). The measurement results were imported and analyzed with R (R Core Team, 2017). All values reported in the paper represent the ratio of the mean fluorescence intensity in the foci of interest over the mean fluorescence intensity within the nucleus. They were computed as follows: (Signal_foci_/Volume_foci_)/(Signal_nucleus_/Volume_nucleus_), where Signal_foci_ corresponds to the integrated fluorescence signal intensity inside of the TALE foci and Signal_nucleus_ corresponds to the total nuclear fluorescence signal intensity. A density greater than one indicates that the signal is higher within the foci.

### Super-resolution microscopy and SMLM data analysis

Super-resolution imaging was performed on a home-made set-up comprising a set of cw lasers (Oxxius LBX-4C-405/488/561/638), an Olympus 100x 1.4 NA oil-immersion objective (UPlanSApo 100x/1.4), and an EM-CCD camera with pixel size of 16 μm x 16 μm (Andor iXon 3 897 Ultra). The objective lens combined with a tube lens of 200 mm of focal length provided a total magnification of the system of 111x, and therefore the pixel size on the object plane was 144 nm x 144 nm. Both photoactivation (405 nm) and excitation (561 nm) lasers were focused on the back focal plane (BFP) of the objective via an appropriate dichroic (Semrock Di03-R561). The position of the focal spot in the BFP in respect to the objective axis was chosen to provide a highly-inclined illumination (HILO) (Tokunaga *et al*, 2008) in order to improve the signal-to-noise-ratio. Fluorescence emission light collected by the objective was filtered at a wavelength centered at 609 nm (Semrock FF01-609/54) before forming the image on the camera. A typical acquisition consisted of 20 000 frames at an integration time of 30 ms.

Raw movies were first analyzed to retrieve the super-localized positions of the single emitters with the ImageJ plugin ThunderSTORM (Ovesný *et al*, 2014), performing a Gaussian fit of the point spread function (PSF) of the recorded single-molecule events. The data was then corrected for sample drift and merging consecutive detections of the same molecule. With the localized emitter positions data, we then generated super-resolved images on a canvas with a pixel size ten-fold increase in respect to the original image. The image was formed by the addition of 2D Gaussian functions of unitary amplitude and size related to their localization accuracy in the retrieved positions.

In order to estimate and compare the size of the TALE clusters, we cropped regions of interest (ROIs) around the clusters in the reconstructed images and normalized them to their maximum intensity pixel value. This was done in order to correct for variability in cell transfection and fluorophore photoactivation, and ensure that only the size of the cluster is accounted for and not the density of detected events in a particular cluster. These ROIs were then segmented and the area of the ROI above an arbitrary intensity threshold was accounted for.

## Acknowledgements

The authors thank Jean-Paul Concordet, Carine Giovannangeli, Marine Charpentier and Anne De Cian from the TACGENE platform for the synthesis of TALE plasmids. We thank Claire Francastel and Jean-Baptiste Boulé for helpful discussion and Loïc Ponger for advises on statistical analysis. This work was supported by funds from the Institut National de la Santé et de la Recherche Médicale, the Centre National de la Recherche Scientifique and the Muséum National d’Histoire Naturelle. SD was supported by a graduate student fellowship from the « Ministère de l’Éductaion Nationale, de l’Enseignement Supérieur et de la Recherche ». LC was supported by the International PhD Program from Institut Curie.

## Author contributions

SD, CE and JL designed the experiments. SD performed the experiments with classical microscopy and analyzed images with the assistance of FL who also provided technical support for TANGO software. LC, KA and II performed PALM microscopy and analyzed images. SD, II, JM, CE and JL analyzed the data and wrote the manuscript.

---

**Supplemental Table 1:** Chromosome instability upon removal of chromosome 7 pericentromeric H3K9me3. Number of nuclei containing either 4 chromosomes 7 and 11 or a different number. Nuclei containing more or less than 4 chromosomes are considered products of aberrant mitosis and proof of genomic instability. Cells grown with doxycycline for 48h (+) are expressing either the TALE-demethylase or the TALE-GFP (control), while cells grown without (−) do not express any construct.

| Doxycycline | TALE-demethylase |  |  |  | TALE-GFP |  |  |  |
| --- | --- | --- | --- | --- | --- | --- | --- | --- |
|  | Chr. 7 |  | Chr. 11 |  | Chr. 7 |  | Chr. 11 |  |
|  | + | - | + | - | + | - | + | - |
| Nuclei with 4 chr. | 1126<br>(93.60%) | 1105<br>(96.09%) | 1169<br>(97.17%) | 1123<br>(97.65%) | 959<br>(98.16%) | 1069<br>(98.25%) | 954<br>(97.65%) | 1068<br>(98.16%) |
| Nuclei with +/- than 4 chr. | 77<br>(6.40%) | 45<br>(3.91%) | 34<br>(2.83%) | 27<br>(2.35%) | 18<br>(1.84%) | 19<br>(1.75%) | 23<br>(2.35%) | 20<br>(1.84%) |
| **Total** | 1203 | 1150 | 1203 | 1150 | 977 | 1088 | 977 | 1088 |

**Supplemental Table 2:** Plasmids used in this study.

| Name | Promoter | Insert | Size (bp) |
| --- | --- | --- | --- |
| **Transient transfections** |  |  |  |
| pJL170 | CMV | FLAG-TALE 394 | 5973 |
| pJL179 | CMV | HA-TALE 394 + GFP | 6183 |
| pJL193 | CMV | HA-TALE 394 + hJMJD2B | 8763 |
| pJL195 | CMV | HA-TALE394 + hJMJD2B-H188A | 8763 |
| pJL203 | CMV | HA-TALE 394 + hJMJD2B + Dendra2 | 9501 |
| pJL206 | CMV | HA-TALE394 + hJMJD2B-H188A + Dendra2 | 9501 |
| **Stable cell lines** |  |  |  |
| pJL204 | pTRE3G | HA-TALE 394 + hJMJD2B | 8560 |
| pJL227 | pTRE3G | HA-TALE 394 + GFP | 5999 |

## References

Ainsztein AM, Kandels-Lewis SE, Mackay AM & Earnshaw WC (1998) INCENP centromere and spindle targeting: Identification of essential conserved motifs and involvement of heterochromatin protein HP1. J Cell Biol 143: 1763–1774

Allshire RC & Madhani HD (2018) Ten principles of heterochromatin formation and function. Nat Rev Mol Cell Biol 19: 229–244

Becker JS, Nicetto D & Zaret KS (2016) H3K9me3-Dependent Heterochromatin: Barrier to Cell Fate Changes. Trends Genet 32: 29–41

Carmena M, Wheelock M, Funabiki H & Earnshaw WC (2012) The chromosomal passenger complex (CPC): from easy rider to the godfather of mitosis. Nat Rev Mol Cell Biol 13: 789–803

Dimitrova E, Turberfield AH & Klose RJ (2015) Histone demethylases in chromatin biology and beyond. EMBO Rep 16: 1620–1639

Erdel F, Rademacher A, Vlijm R, Tünnermann J, Frank L, Weinmann R, Schweigert E, Yserentant K, Hummert J, Bauer C, et al (2020) Mouse Heterochromatin Adopts Digital Compaction States without Showing Hallmarks of HP1-Driven Liquid-Liquid Phase Separation. Mol Cell 78: 236–249.e7

Fernandez Garcia M, Moore CD, Schulz KN, Alberto O, Donague G, Harrison MM, Zhu H & Zaret KS (2019) Structural Features of Transcription Factors Associating with Nucleosome Binding. Mol Cell 75: 921–932

Fukagawa T & Earnshaw WC (2014) The centromere: Chromatin foundation for the kinetochore machinery. Dev Cell 30: 496–508

Funabiki H (2019) Correcting aberrant kinetochore microtubule attachments: a hidden regulation of Aurora B on microtubules. Curr Opin Cell Biol 58: 34–41

Giet R & Glover DM (2001) Drosophila aurora B kinase is required for histone H3 phosphorylation and condensin recruitment during chromosome condensation and to organize the central spindle during cytokinesis. J Cell Biol 152: 669–681

Gordon GS & Wright A (2000) DNA segregation in bacteria. Annu Rev Microbiol 54: 681–708

Hayashi-Takanaka Y, Yamagata K, Nozaki N & Kimura H (2009) Visualizing histone modifications in living cells: Spatiotemporal dynamics of H3 phosphorylation during interphase. J Cell Biol 187: 781–790

Hayden KE (2012) Human centromere genomics: now it’s personal. Chromosom Res 20: 621–33

Hayden KE, Strome ED, Merrett SL, Lee H-R, Rudd MK & Willard HF (2013) Sequences Associated with Centromere Competency in the Human Genome. Mol Cell Biol 33: 763–772

Jankele R & Svoboda P (2014) TAL effectors: Tools for DNATargeting. Brief Funct Genomics 13: 409–419

Janssen A, Colmenares SU & Karpen GH (2018) Heterochromatin: Guardian of the Genome. Annu Rev Cell Dev Biol 34: 265–288

Klemm SL, Shipony Z & Greenleaf WJ (2019) Chromatin accessibility and the regulatory epigenome. Nat Rev Genet 20: 207–220

Kruitwagen T, Denoth-Lippuner A, Wilkins BJ, Neumann H & Barral Y (2015) Axial contraction and short-range compaction of chromatin synergistically promote mitotic chromosome condensation. Elife 4: e10396

Kumar A & Kono H (2020) Heterochromatin protein 1 (HP1): interactions with itself and chromatin components. Biophys Rev 12: 387–400

Larson AG, Elnatan D, Keenen MM, Trnka MJ, Johnston JB, Burlingame AL, Agard DA, Redding S & Narlikar GJ (2017) Liquid droplet formation by HP1α suggests a role for phase separation in heterochromatin. Nature 547: 236–240

Lavoie BD, Hogan E & Koshland D (2004) In vivo requirements for rDNA chromosome condensation reveal two cell-cycle-regulated pathways for mitotic chromosome folding. Genes Dev 18: 76–87

Liu X, Song Z, Huo Y, Zhang J, Zhu T, Wang J, Zhao X, Aikhionbare F, Zhang J, Duan H, et al (2014) Chromatin protein HP1α interacts with the mitotic regulator borealin protein and specifies the centromere localization of the chromosomal passenger complex. J Biol Chem 289: 20638–20649

Mahlke MA & Nechemia-Arbely Y (2020) Guarding the genome: Cenp-a-chromatin in health and cancer. Genes 11: 810

Martins NMC, Cisneros-Soberanis F, Pesenti E, Kochanova NY, Shang WH, Hori T, Nagase T, Kimura H, Larionov V, Masumoto H, et al (2020) H3K9me3 maintenance on a human artificial chromosome is required for segregation but not centromere epigenetic memory. J Cell Sci 133: jsc242610

McManus KJ, Biron VL, Heit R, Underhill DA & Hendzel MJ (2006) Dynamic changes in histone H3 lysine 9 methylations: Identification of a mitosis-specific function for dynamic methylation in chromosome congression and segregation. J Biol Chem 281: 8888–8897

McNulty SM & Sullivan BA (2018) Alpha satellite DNA biology: finding function in the recesses of the genome. Chromosom Res 26: 115–138

Molina O, Carmena M, Maudlin IE & Earnshaw WC (2016) PREditOR: a synthetic biology approach to removing heterochromatin from cells. Chromosom Res 24: 495–509

Moscou MJ & Bogdanove AJ (2009) Recognition by TAL Effectors. Science 326: 1501

Mozzetta C, Boyarchuk E, Pontis J & Ait-Si-Ali S (2015) Sound of silence: The properties and functions of repressive Lys methyltransferases. Nat Rev Mol Cell Biol 16: 499–513

Muller H, Gil J & Drinnenberg IA (2019) The Impact of Centromeres on Spatial Genome Architecture. Trends Genet 35: 565–578

Müller S & Almouzni G (2017) Chromatin dynamics during the cell cycle at centromeres. Nat Rev Genet 18: 192–208

Ninova M, Tóth KF & Aravin AA (2019) The control of gene expression and cell identity by H3K9 trimethylation. Development 146: dev181180

Nishibuchi G & Déjardin J (2017) The molecular basis of the organization of repetitive DNA-containing constitutive heterochromatin in mammals. Chromosom Res 25: 77–87

Nishibuchi G & Nakayama JI (2014) Biochemical and structural properties of heterochromatin protein 1: Understanding its role in chromatin assembly. J Biochem 156: 11–20

Ohishi T, Muramatsu Y, Yoshida H & Seimiya H (2014) TRF1 Ensures the Centromeric Function of Aurora-B and Proper Chromosome Segregation. Mol Cell Biol 34: 2464–2478

Ohzeki J, Larionov V, Earnshaw WC & Masumoto H (2019) De novo formation and epigenetic maintenance of centromere chromatin. Curr Opin Cell Biol 58: 15–25

Ollion J, Cochennec J, Loll F, Escudé C & Boudier T (2013) TANGO: A generic tool for high-throughput 3D image analysis for studying nuclear organization. Bioinformatics 29: 1840–1841

Ollion J, Cochennec J, Loll F, Escudé C & Boudier T (2015a) Analysis of nuclear organization with TANGO, Software for high-throughput quantitative analysis of 3D fluorescence microscopy images. Methods Mol Biol 1228: 203–222

Ollion J, Loll F, Cochennec J, Boudier T & Escudé C (2015b) Proliferation-dependent positioning of individual centromeres in the interphase nucleus of human lymphoblastoid cell lines. Mol Biol Cell 26: 2550–60

Ovesný M, Křížek P, Borkovec J, Švindrych Z & Hagen GM (2014) ThunderSTORM: A comprehensive ImageJ plug-in for PALM and STORM data analysis and super-resolution imaging. Bioinformatics 30: 2389–2390

Peters AHFM, Carroll O, Scherthan H, Mechtler K, Sauer S, Scho C, Weipoltshammer K, Pagani M, Lachner M, Kohlmaier A, et al (2001) Loss of the Suv39h Histone Methyltransferases Impairs Mammalian Heterochromatin and Genome Stability. Cell 107: 323–337

Petrova B, Dehler S, Kruitwagen T, Heriche J-K, Miura K & Haering CH (2013) Quantitative Analysis of Chromosome Condensation in Fission Yeast. Mol Cell Biol 33: 984–998

Pluta AF, Cooke CA & Earnshaw WC (1990) Structure of the human centromere at metaphase. Trends Biochem Sci 15: 181–185

R Core Team (2017) R: A Language and Environment for Statistical Computing. R Found Stat Comput Vienna, Austria: https://www.R-project.org/

Ruppert JG, Samejima K, Platani M, Molina O, Kimura H, Jeyaprakash AA, Ohta S & Earnshaw WC (2018) HP 1α targets the chromosomal passenger complex for activation at heterochromatin before mitotic entry. EMBO J 37: e97677

Sales-Gil R & Vagnarelli P (2020) How HP1 Post-Translational Modifications Regulate Heterochromatin Formation and Maintenance. Cells 9: 1460

Schalch T & Steiner FA (2017) Structure of centromere chromatin: from nucleosome to chromosomal architecture. Chromosoma 126: 443–455

Schumacher MA, Tonthat NK, Lee J, Rodriguez-Castañeda FA, Chinnam NB, Kalliomaa-Sanford AK, Ng IW, Barge MT, Shaw PLR & Barillà D (2015) Structures of archaeal DNA segregation machinery reveal bacterial and eukaryotic linkages. Science 349: 1120–1124

Sharma AB, Dimitrov S, Hamiche A & Van Dyck E (2019) Centromeric and ectopic assembly of CENP-A chromatin in health and cancer: old marks and new tracks. Nucleic Acids Res 47: 1051–1069

Slee RB, Steiner CM, Herbert B-S, Vance GH, Hickey RJ, Schwarz T, Christan S, Radovich M, Schneider BP, Schindelhauer D, et al (2012) Cancer-associated alteration of pericentromeric heterochromatin may contribute to chromosome instability. Oncogene 31: 3244–53

Stellfox ME, Bailey AO & Foltz DR (2013) Putting CENP-A in its place. Cell Mol Life Sci 70: 387–406

Strom AR, Emelyanov AV, Mir M, Fyodorov DV, Darzacq X & Karpen GH (2017) Phase separation drives heterochromatin domain formation. Nature 547: 241–245

Tokunaga M, Imamoto N & Sakata-Sogawa K (2008) Highly inclined thin illumination enables clear single-molecule imaging in cells. Nat Methods 5: 159–161

Voytas DF & Joung JK (2009) DNA Binding Made Easy. Science 326: 1491–1492

Wei Y, Yu L, Bowen J, Gorovsky MA & David Allis C (1999) Phosphorylation of histone H3 is required for proper chromosome condensation and segregation. Cell 97: 99–109

Whetstine JR, Nottke A, Lan F, Huarte M, Smolikov S, Chen Z, Spooner E, Li E, Zhang G, Colaiacovo M, et al (2006) Reversal of Histone Lysine Trimethylation by the JMJD2 Family of Histone Demethylases. Cell 125: 467–481

Willard HF (1985) Chromosome-specific organization of human alpha satellite DNA. Am J Hum Genet 37: 524–532

Yi Q, Chen Q, Liang C, Yan H, Zhang Z, Xiang X, Zhang M, Qi F, Zhou L & Wang F (2018) HP 1 links centromeric heterochromatin to centromere cohesion in mammals. EMBO Rep 19: e45484

